# Global gradients in intertidal species richness and functional groups

**DOI:** 10.1101/2020.12.16.423020

**Authors:** Jakob Thyrring, Lloyd S. Peck

## Abstract

Whether global latitudinal diversity gradients exist in rocky intertidal α-diversity and across functional groups remains unknown. Using literature data from 433 intertidal sites, we investigated α-diversity patterns across 155° of latitude, and whether local-scale or global-scale structuring processes control α-diversity. We, furthermore, investigated how the relative composition of functional groups (algae, grazers, predators and suspension-feeders) changes with latitude. α-diversity differed among hemispheres with a mid-latitudinal peak in the north, and a non-significant unimodal pattern in the south, but there was no support for a latitudinal diversity gradient. Although global-scale drivers had no discernible effect, the local-scale drivers significantly affected α-diversity, and our results reveal that latitudinal diversity gradients are outweighed by local-processes. In three functional groups: predators, grazers and suspension-feeders diversity declined with latitude, coinciding with an inverse gradient in algae. Overall, we propose more studies are needed on the magnitude and influence of physical and biotic drivers across multiple scales.

## Introduction

The latitudinal diversity gradient in species richness across ecosystems and various functional groups has been a major research topic that has intrigued scientists since at least Darwin (1859) and Wallace (1878). Over time, many hypotheses have been proposed to explain this seemingly general ecological pattern in marine and terrestrial ecosystems. Ecological and physical hypotheses dominate the discussions and include drivers such as habitat area, stability, speciation rates, energy availability, and temperature [3–5]. However, the geographic, functional and taxonomic generality of latitudinal diversity gradients remain lively debated as unimodal, bimodal and inverse gradients emerge across clades, habitats and latitudes [6–9]. In the marine realm, latitudinal gradients in some groups have been shown to be closely related to oceanographic covariates, such as water temperature [10], yet mammal richness peaks at high latitudes [11], and more species are found in polar soft-sediment habitats than at many lower latitudes [12].

While gradients in species diversity have received most attention, latitudinal changes across different functional groups (evaluated here by food acquisition, see Materials and Methods) remain less studied despite their importance for ecosystem functioning; Macroalgal fronds create protective microhabitats increasing biodiversity and sheltering understory species from environmental stress [13,14], suspension-feeders are important benthic-pelagic energy couplers [15], and predation is widely accepted as a central structuring process in the composition and abundance of species [16,17]. Latitudinal studies have shown suspension-feeders dominate benthic systems in fully marine environments at high latitudes [15,18], while diversity of coastal macroalgae peaks at mid-latitudes [19], and predation pressure decreases with latitude and depths from shallow shelves to deep oceans [20,21]. Most functional group studies have focused on a narrow set of taxa, and global-scale assembly-wide investigations encompassing both hemispheres are rare. Thus, studies demonstrating global patterns in various functional groups are needed to understand spatial patterns, biological interactions and ecosystem resilience to climate change.

Intertidal shores rank as one of the most studied marine habitats, and are often seen as harbingers for the effects of climate change and invasive species [22]. Regional-scale intertidal studies have found richness gradients of gastropods along coastlines in the eastern Pacific Ocean [23,24]. However, no latitudinal diversity gradient of gastropods [25] or macroalgal [26] was found on a global-scale across oceans, and assembly-wide studies have found missing [27–29] or inverse [30] gradients. Conflicting and missing gradients suggests that richness is determined by regional or local (and not global) scale processes through biological interactions and small-scale overlapping environmental gradients. For example, high habitat heterogeneity creates environmental stress mosaics that are more important in shaping biological patterns than latitudinal environmental gradients [31,32], and different oceanographic covariates measured on latitudinal and longitudinal scales across Europe had limited effect on local species richness [33]. However, the large scale intertidal studies necessary to evaluate global scale patterns and processes are generally missing, but see [25,28].

Global assessments of intertidal biodiversity have been hindered by inadequate estimates of intertidal areas [34], and data scarcity from polar intertidal shores. However, in recent decades intertidal diversity data from both polar regions have advanced [30,35–39], providing novel opportunities to study biodiversity patterns and variation in specific functional groups on a worldwide scale. In this study, we examined latitudinal diversity gradients in local intertidal species richness (termed α-diversity) and four functional groups from 433 locations throughout the northern and southern hemisphere, from the high Arctic to the Antarctic. Intertidal ecosystems are comprised of a variety of habitats ranging from tidal flats to mangrove forests, each supporting unique communities and taxa. Therefore, to control for habitat effects, we specifically focus on rocky shores and boulderfields, as they represent the most studied intertidal habitats and are present in all oceans. We aim to examine patterns in α-diversity and functional groups across 155° of latitude, and test the following intertidal hypotheses: (1) α-diversity decreases with increasing latitude, as evident in other ecosystems; (2) biodiversity is controlled by local-scale structuring processes rather than global-scale oceanographic features.

## Results

### Latitudinal α-diversity gradients

We extracted data from 433 intertidal sites between 74.8°S and 80.5°N (Fig. 1). Only 10 sites were located at latitudes above 70°, south and north combined. Generalized linear mixed models (GLMM) indicated non-significant relationships among α-diversity and latitude (Fig. 2; Table 1), but the latitudinal patterns were different among hemispheres; In the southern hemisphere, α-diversity displayed a non-significant unimodal pattern with a possible slight decline at the highest latitude (Fig. 2; Table 1), while northern hemisphere α-diversity peaked at mid-latitudes between 30–40°N (Fig 2; Table 1).

**Figure 1:**
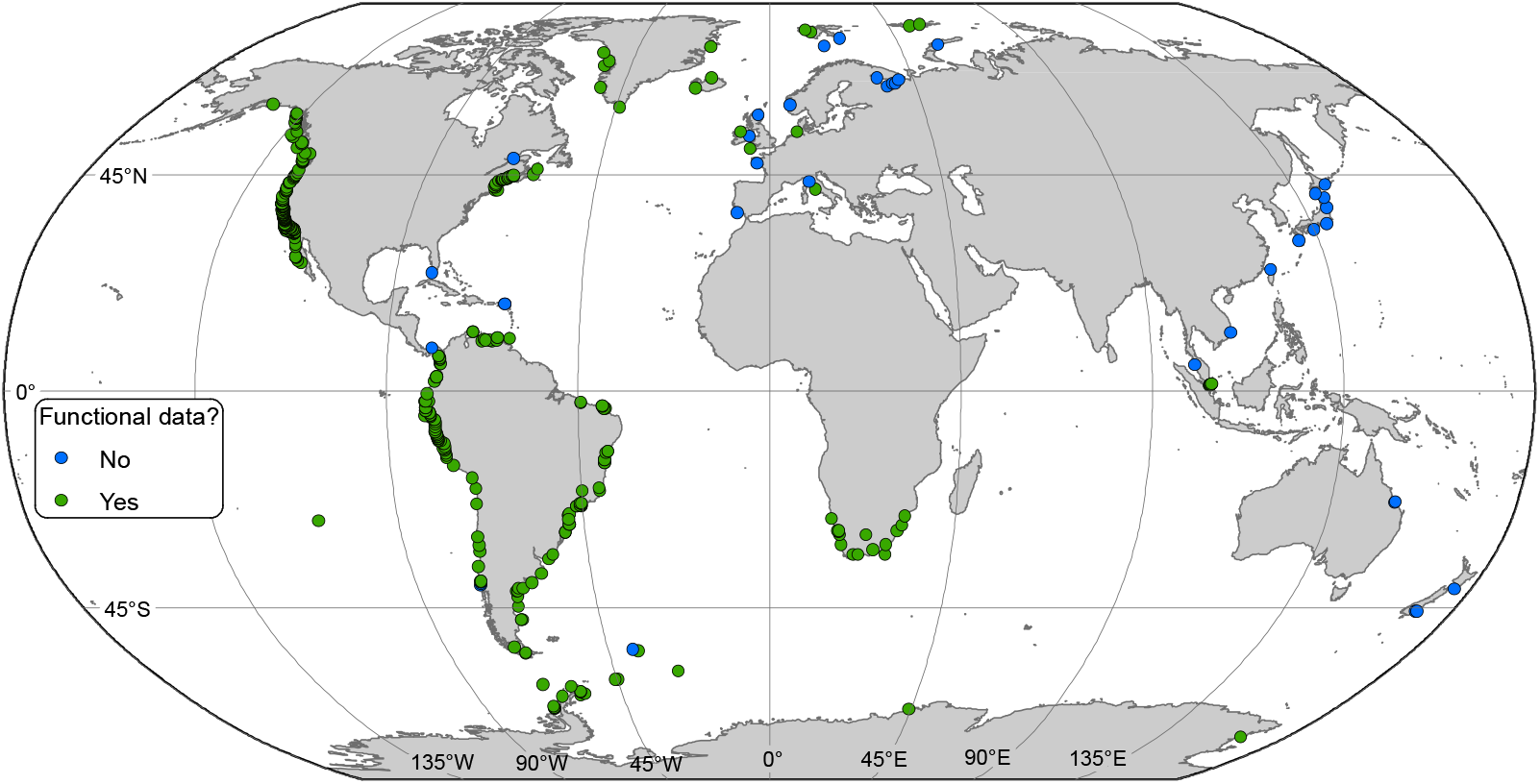
Locations of rocky intertidal sampling sites. Some sites are not visible because close proximity. Functional diversity data were available in green sites.

**Figure 2:**
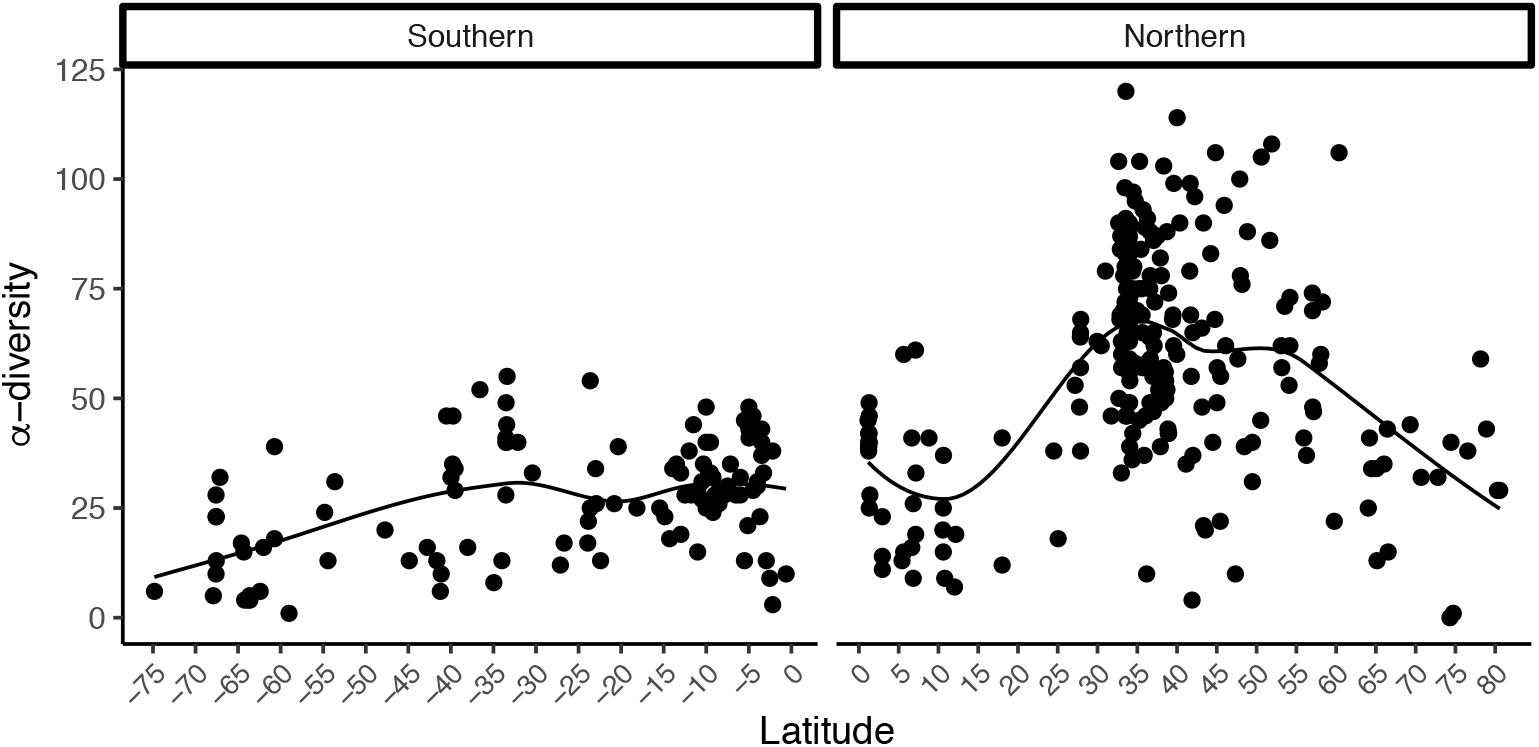
Latitudinal patterns in rocky intertidal α-diversity plotted against latitude. Data are split into southern and northern hemisphere. R^2^ values and a best-fit locally weighted scatterplot smoother (LOESS) was added to aid visual interpretation.

**Table 1:**
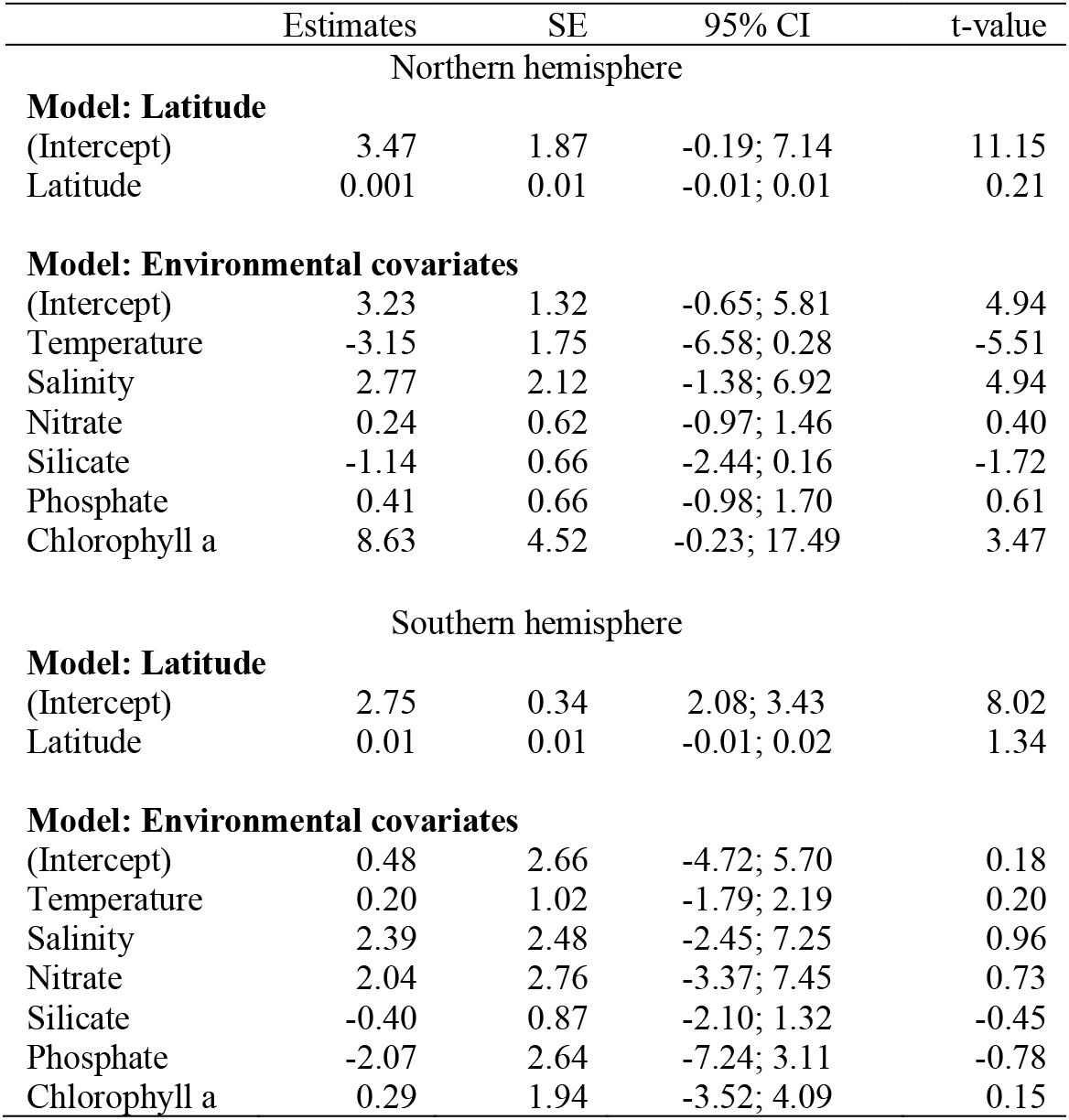
Latitudinal-scale generalized linear mixed model summaries. Estimated parameters, standard error (SE), bootstrapped 95% confidence intervals (95% CI) and p-values are reported for the relationship between α-diversity and environmental covariates.

|  | Estimates | SE | 95% CI | t-value |
| --- | --- | --- | --- | --- |
| Northern hemisphere |  |  |  |  |
| **Model: Latitude** |  |  |  |  |
| (Intercept) | 3.47 | 1.87 | -0.19; 7.14 | 11.15 |
| Latitude | 0.001 | 0.01 | -0.01; 0.01 | 0.21 |
| **Model: Environmental covariates** |  |  |  |  |
| (Intercept) | 3.23 | 1.32 | -0.65; 5.81 | 4.94 |
| Temperature | -3.15 | 1.75 | -6.58; 0.28 | -5.51 |
| Salinity | 2.77 | 2.12 | -1.38; 6.92 | 4.94 |
| Nitrate | 0.24 | 0.62 | -0.97; 1.46 | 0.40 |
| Silicate | -1.14 | 0.66 | -2.44; 0.16 | -1.72 |
| Phosphate | 0.41 | 0.66 | -0.98; 1.70 | 0.61 |
| Chlorophyll a | 8.63 | 4.52 | -0.23; 17.49 | 3.47 |
| Southern hemisphere |  |  |  |  |
| **Model: Latitude** |  |  |  |  |
| (Intercept) | 2.75 | 0.34 | 2.08; 3.43 | 8.02 |
| Latitude | 0.01 | 0.01 | -0.01; 0.02 | 1.34 |
| **Model: Environmental covariates** |  |  |  |  |
| (Intercept) | 0.48 | 2.66 | -4.72; 5.70 | 0.18 |
| Temperature | 0.20 | 1.02 | -1.79; 2.19 | 0.20 |
| Salinity | 2.39 | 2.48 | -2.45; 7.25 | 0.96 |
| Nitrate | 2.04 | 2.76 | -3.37; 7.45 | 0.73 |
| Silicate | -0.40 | 0.87 | -2.10; 1.32 | -0.45 |
| Phosphate | -2.07 | 2.64 | -7.24; 3.11 | -0.78 |
| Chlorophyll a | 0.29 | 1.94 | -3.52; 4.09 | 0.15 |

In both hemispheres, GLMMs failed to detect significant effects for latitude (Table 1), or for six latitudinal-scale oceanographic covariates on α-diversity (Fig. 3; Table 1). On the contrary, localscale models revealed significant relationships between α-diversity and the four local-scale covariates ice scour, macroalgal cover, salinity and wave exposure (Fig. 4, Table 2). A positive relationship was found between α-diversity and salinity and macroalgal cover (Fig 5a,c), with salinity having the strongest effect (R^2^ = 0.19, 95% CI = 4.82 – 8.52, Fig. 4). Wave exposure and ice scour had negative effects on α-diversity (Fig. 5b,d). The non-linear relationship between wave exposure and α-diversity showed that high levels of wave exposure were necessary to affect levels of α-diversity (Fig. 5b). Fewer species inhabited ice-scoured shores compared to ice-free shores (95% CI = −0.57 – −0.23, Table 2), although the model only explained around 5% (R^2^ = 0.05) of the variation due to large differences in the number of species on both ice-scoured and ice-free shores (Fig. 5c).

**Figure 3:**
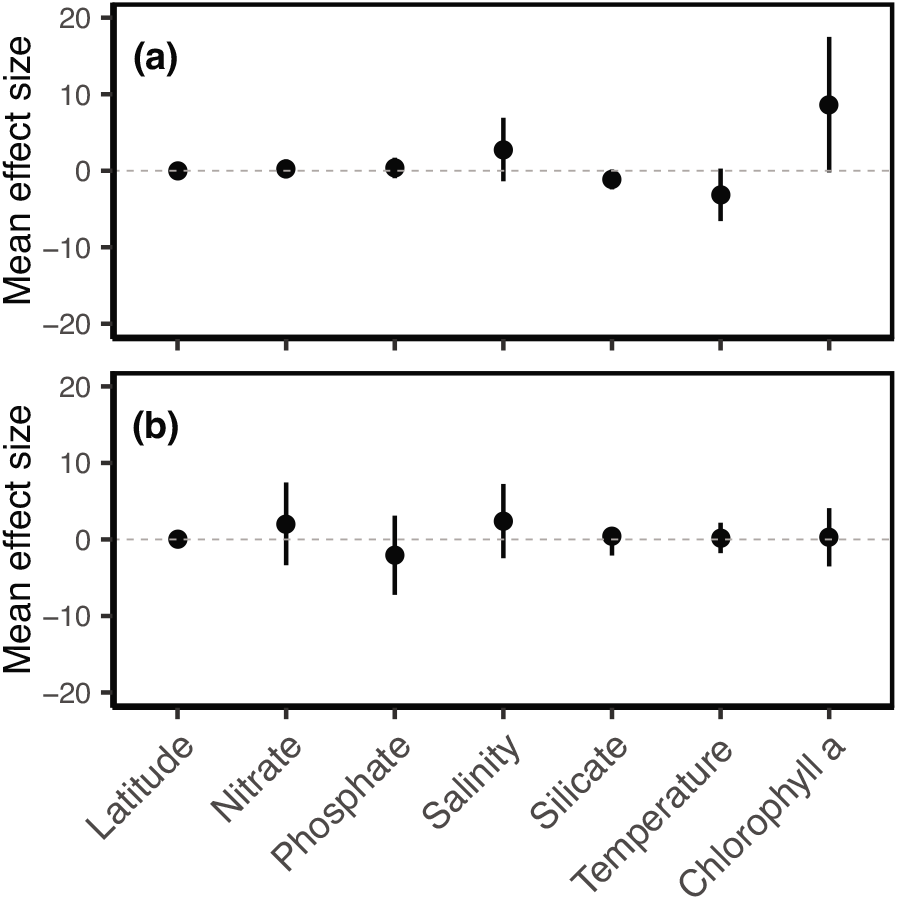
Latitudinal-scale mean effect sizes and direction of latitude, nitrate, phosphate, salinity, silicate, sea surface temperature, and chlorophyll *a* on α-diversity in **(a)** northern and **(b)** southern hemisphere. Non-significance of regression parameters is identified as bootstrapped 95% confidence intervals (error bars) crossing zero.

**Figure 4:**
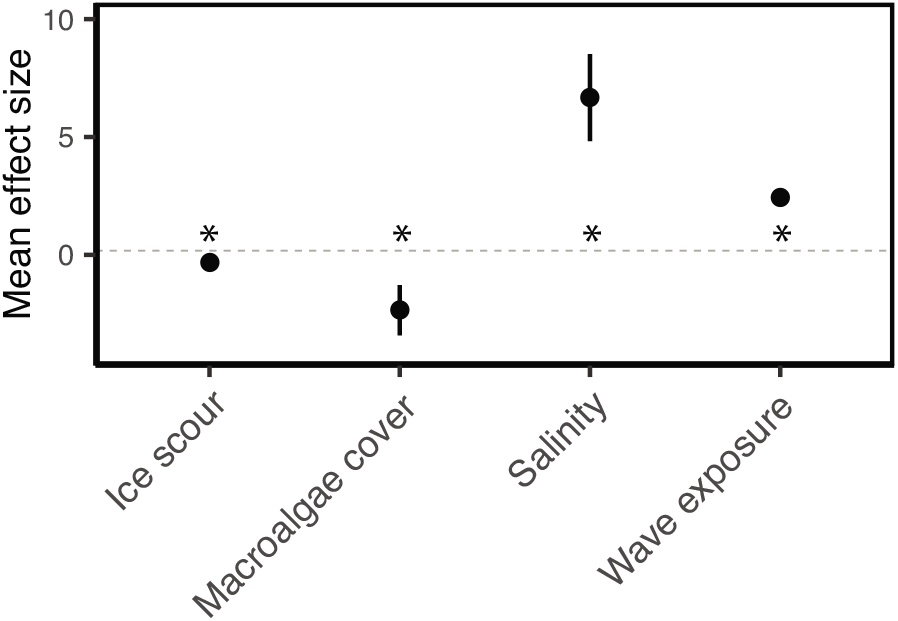
Local-scale mean effect sizes and direction of ice scour, macroalgal cover, salinity, and wave exposure on α-diversity estimated from individual models. Significance of regression parameters is identified as bootstrapped 95% confidence intervals (error bars) not crossing zero (* indicate significance).

**Table 2:** Local-scale model summaries with individual models and distributions indicated for each model. Estimated parameters, standard error (SE), bootstrapped 95% confidence intervals (95% CI), z-values and p-values are reported for the relationship between α-diversity and environmental covariates.

|  |  |  |  |  |  |
| --- | --- | --- | --- | --- | --- |
| Salinity $R^2 = 0.19$ (GLM) | | | | | |
|  | Estimates | SE | 95% CI | z-value | p-values |
| (Intercept) | -3.201 | 0.891 | -4.94; -1.47 | -3.6 | <0.0001 |
| Salinity | 6.676 | 0.946 | 4.82; 8.52 | 7.06 | <0.0001 |
| Wave exposure (GAM) |  |  |  |  |  |
| $R^2 = 0.12$ | | | | | |
|  | Estimates | SE | 95% CI | z-value | p-values |
| (Intercept)* | 2.435 | 0.029 | 2.38; 2.43 | 82.4 | <0.0001 |
| Ice scour (GLM) |  |  |  |  |  |
| $R^2 = 0.05$ | | | | | |
|  | Estimates | SE | 95% CI | z-value | p-values |
| (Intercept) | 2.353 | 0.055 | 2.24; 2.46 | 42.7 | <0.0001 |
| Ice Scour | -0.401 | 0.087 | -0.57; -0.23 | -4.59 | <0.0001 |
| Macroalgal cover (GLM) |  |  |  |  |  |
| $R^2 = 0.14$ | | | | | |
|  | Estimates | SE | 95% CI | z-value | p-values |
| (Intercept) | 6.133 | 0.386 | 5.37; 6.89 | 15.9 | <0.0001 |
| Cover | -2.35 | 0.547 | -3.42; -1.28 | -4.3 | <0.0001 |

\*Approximate significance of locally weighted scatterplot wave exposure smoother (LOESS): Estimated df = 8.40, p-value < 0.0001

**Figure 5:**
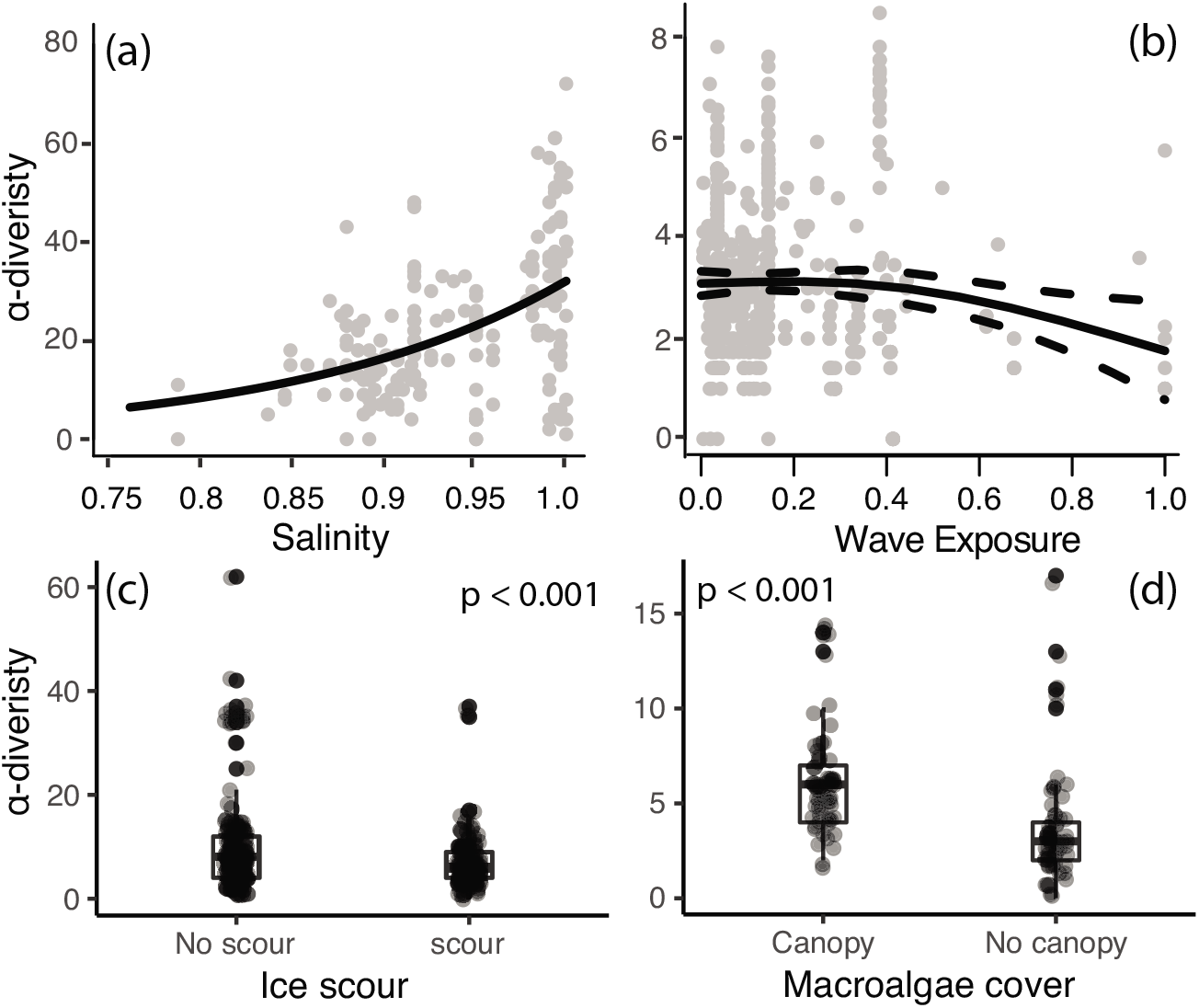
Local-scale relationships between **(a)** salinity, **(b)** wave exposure **(c)** ice scour and **(d)** macroalgae cover on α-diversity. A best-fit locally weighted scatterplot smoother (LOESS) (panel **a,b**) and boxplots (panel **c,d**) has been added to aid visual interpretation.

### Latitudinal gradients in functional groups

Four studied functional groups (algae, grazers, predators, suspension-feeders) displayed different latitudinal patterns (Fig. 6). Predators, grazers and suspension-feeders declined with latitude in both hemispheres, coinciding with an increase in algal richness (Fig. 6). In general, the relative distribution of the functional groups changed faster across latitudes in the southern hemisphere where rapid changes were observed around 20°S, while northern hemisphere communities were more consistent from 25–60°N (Fig. 6). We were unable to describe functional groups at latitudes 15–25°N as no data were available (indicated as white bars in figure 6).

**Figure 6:**
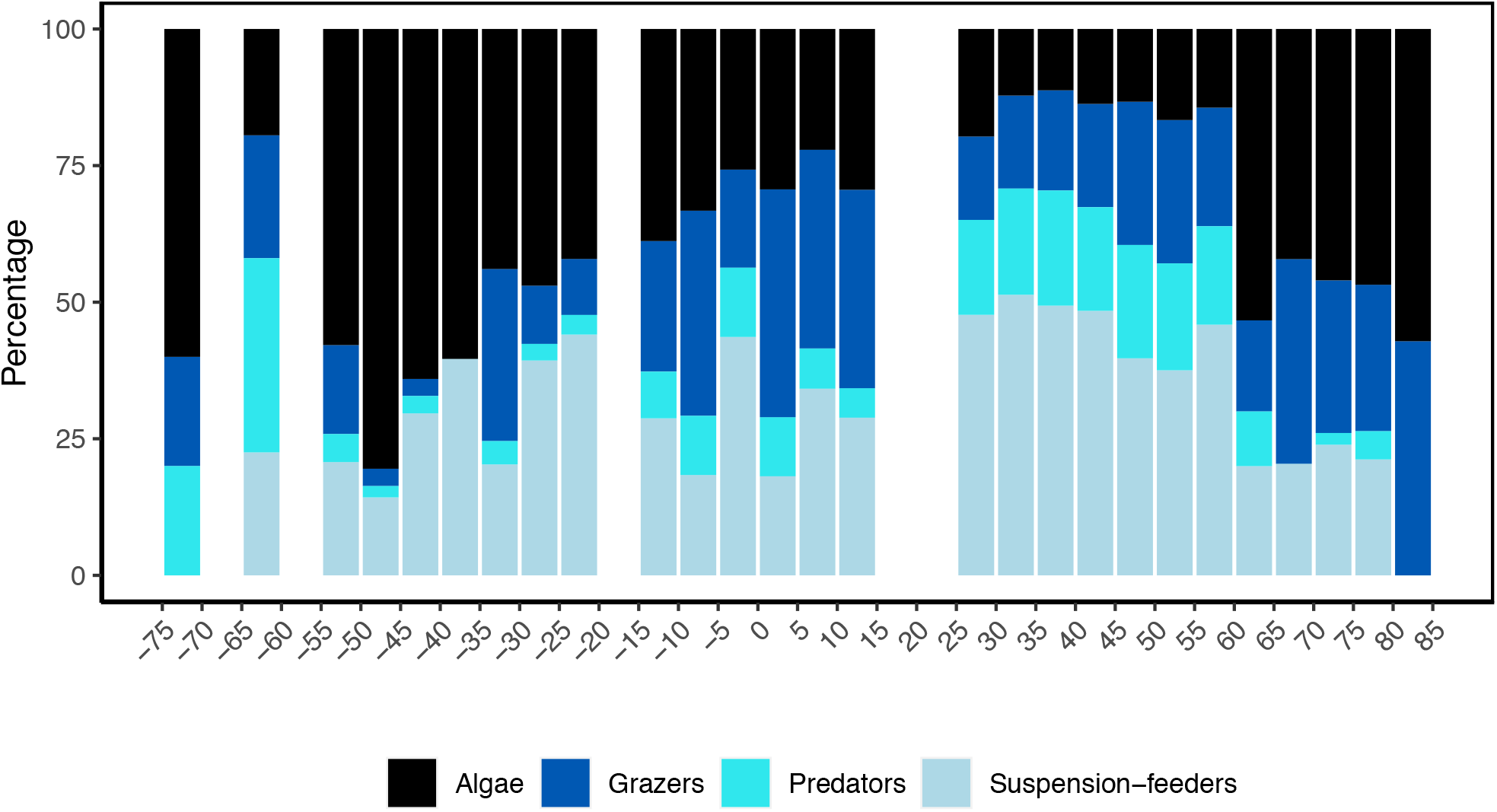
Latitudinal variation in the relative number of species from four functional groups in the rocky mid-intertidal between 65°S to 85°N, where negative values denote southern hemisphere latitudes. No data were available within the 5° latitudinal bands where bars are missing.

In the southern hemisphere the proportion of predators and grazers declined beyond around 20°S, together forming 6.9% (±6.6 s.d) of species found between 20–55°S, but constituting 18.3% (±15.1 s.d) of species in communities at latitudes less than 20°S (Fig. 6). Suspension-feeders were common in low-to mid-latitudes, constituting 34.6% (±9.6 s.d) of the species present between 20–45°S (Fig. 6). The relative proportion of algal species increased with latitude, with a small decline at sites between 60–65° (Fig. 6). In the northern hemisphere, predators were abundant at low and midlatitudes, accounting on average for 18.2% (±5.8 s.d) of the populations from 0-60°N, before declining to 4.7 % (±4.1 s.d) at high latitudes between 61–80°N. The number of grazers decreased markedly from around 15°N, but stabilized at 25.4% (±6.4 s.d) across 25°-80°N (Fig. 6). Suspensionfeeders were the dominant functional group on shores in the northern hemisphere, especially between 25–60°N, but decreased to 21.4% (±1.8 s.d) at high latitudes between 60–80°N (Fig. 6). Algal species was the most dominant group at high latitudes (Fig. 6). In both hemispheres, the community composition at the highest latitudes was substantially different from elsewhere; on these high polar shores suspension feeders were absent, and only a few predators were present, while algae dominated.

## Discussion

### Latitudinal α-diversity gradient

Latitudinal diversity gradients have been recognized for centuries, and the generality of this phenomenon has been investigated ever since. Recent state-of-the-art meta-analyses have shown support for latitudinal gradients in various marine and terrestrial ecosystems [6,40], and latitudinal gradients in assembly composition and biological interactions have also been identified [21,41,42]. Although latitudinal diversity gradients are widely quoted as one of the basic laws in ecology [43], studies investigating these patterns on a global scale have largely overlooked intertidal ecosystems, despite their global range and importance as unique environments. Intertidal studies in the north eastern Pacific Ocean have reported no latitudinal gradients in species richness [27,29,44]. It has been suggested that the latitudinal diversity gradient is biogeographically structured[45], and since the above investigations were conducted in a temperate biogeographic region, a gradients might not occur. For example, while gastropod richness is relatively stable in the temperate region, a steep rise in richness occurs in the tropical region [23]. However, the few published global-scale studies have revealed contradictory latitudinal patterns with richness of macroalgae [26] and small echinoderms peaking at high latitudes, while large echinoderms peak at low latitudes [46]. Consequently, no latitudinal trend in intertidal γ-diversity was found across 13 marine regions, as overall assemblage richness was similar everywhere despite strong species specific patterns [28]. While the studies above benefitted from analysing data collected using a standardized protocol across latitudes, they were limited by only including a smaller number of sampling sites (~70 sites), which may have restricted their capacity to demonstrate latitudinal patterns. Compared to these studies, we expanded the spatial resolution six-fold and present α-diversity data from 433 rocky shore sites across 155° of latitude, yet there was no support for the latitudinal α-diversity gradient hypothesis. Latitudinal patterns were, however, markedly different among hemispheres, with a midlatitude peak in the north and a non-significant unimodal trend in the south. We also demonstrated different functional groups had different gradients (see later discussion).

Northern hemisphere α-diversity displayed a distinct mid-latitude peak, a pattern also reported for open water biodiversity [7]. The cause behind this remains elusive but hypotheses include a mid-dominance effect where high- and low-latitude species ranges overlap [47], and that high temperatures near the equator, and low temperature and ice cover at the poles, may limit diversity outside of the temperate zone. However, we show that atmospheric temperatures do not control intertidal α-diversity, and ice-scour only has a minor impact on the number of species found. The open water mid-latitudinal richness peak has recently been suggested to be an artefact of limited sampling in less studied tropical regions [48]. This asymmetric sampling holds true for the northern hemisphere rocky intertidal ecosystem as well; While we found data from less than 10 sites at latitudes above 70° (south and north combined), 44% (n = 193) of all studies were from mid-latitude rocky shores in Europe and North America between 24–60°N. Data from low-latitude regions is furthermore limited because rocky shorelines are scarce in the tropics [23], and the majority of tropical studies have focused on specific taxonomic groups, and not assembly-wide patterns [e.g. 26,28,48-53], but see [55]. Thus, mid-latitudinal rocky shores, especially in the northern hemisphere, are among the most sampled globally, and future tropical collections may change understanding of biodiversity patterns [55]. This disproportionate sampling pattern is also present in other marine habitats, where sampling intensity is highest at mid-latitudes in the northern hemisphere [48]. Polar data were also limited in the southern hemisphere where species diversity was relatively constant across latitudes with a non-significant dip at the highest latitude. A factor that powerfully affects intertidal diversity in the high polar regions is that most intertidal areas are encased in ice for large parts, if not all of the year, which reduces intertidal biodiversity to zero in these areas. Thus, the Arctic permafrost coastline represent around 34% of the world’s coastline [56], and Antarctica accounts for around 2.7% (45,317 km). However, in the Antarctic only around 12% of that is ice free in summer (5468 km) and at both poles much less, if any, is ice free in winter [35]. This lack of ice-free intertidal areas in high polar sites restricts the capacity for communities to establish, and limits the development of macroalgal species [57–59]. However, high polar shores are seldom studied, and in this study high latitude Antarctic shorelines were only represented by a single site, thus this depression in number of species should be interpreted with caution. In ice-free areas, and at lower Antarctic latitudes, previous biodiversity studies have shown intertidal species richness to increase with latitude from the southern Atlantic ocean to the Antarctic Peninsula [30,60]. The situation is clearly complex and, in general, more information on the influence of geographic sampling biases across latitudes are, together with more data from more sites, especially the polar and tropical regions, required to understand and model large-scale biodiversity patterns.

Latitudinal diversity gradients have been identified in subtidal habitats (e.g. [3,10]), and the contrasting patterns in biodiversity gradients between intertidal and subtidal ecosystems may originate from differences in the factors controlling α-diversity. Subtidal benthic habitats experience relatively homogenous environmental conditions across small to moderate spatial scales, and large-scale latitudinal diversity gradients are therefore less impacted by local factors and reflect larger-scale changes in environmental conditions. For instance, decreasing water temperatures profoundly affect physiological performance of marine ectotherms (e.g. growth, reproduction and activity, muscle performance) and special adaptations are required to live in cold waters [35,61]. Latitudinal biodiversity gradients in ectotherms have, therefore, been attributed to changes in water temperatures in subtidal habitats [3,10,11]. Our data run contrary to this and indicate that in the intertidal there is no effect of latitude, or any of the correlated environmental drivers (e.g. primary production, salinity, water temperature) on α-diversity. Since latitude is a surrogate for environmental factors that co-vary with latitude, other large-scale factors, such as daylength, are likely not to be significant either. α-diversity may display a different latitudinal patterns than γ-diversity as local environmental conditions determine local richness [44,62]. Indeed, we demonstrate significant effects of local-scale variation in salinity, wave exposure, ice scour and biogenic habitats (i.e. macroalgal cover), indicating that α-diversity is determined by small-to regional-scale processes through biological interactions and small-scale overlapping environmental gradients. Accordingly, we suggest that the patterns observed here in intertidal α-diversity arise from intertidal rocky shores being highly heterogenous habitats, that change on small temporal and spatial scales, and these factors outweigh latitudinal drivers of biodiversity. In support of this view, previous studies have shown how unique environmental conditions are created along coastlines by small-scale variations in a range of factors including temperature, pH, surface orientation (i.e. in the northern hemisphere south facing surfaces are hotter than north facing). Even wave splashing and low water timing (i.e. low tides occurring in the tropics during the hottest time of the day are more harmful than low tides in the middle of the night [22]) have been shown to be important. All these factors interact and change in non-latitudinally related patterns [31,63,64]. Biogenic factors, especially macroalgal cover, can create intertidal microhabitats sheltering organisms from extreme conditions and increase α-diversity (see Figure 5). Therefore, local environmental conditions and microhabitats can, in combination, functionally override latitudinal stress or energy gradients. Thus, although the strength and direction of environmental stressors changes across latitudes (i.e. from ice scour and extreme sub-zero temperature in polar regions to desiccation and acute heat stress in the tropics), the combined stress experienced by resident populations may roughly balance out across most latitudes and produce no clear gradient. A conclusion from this is that instead of focusing on large-scale drivers that implicitly assumed patterns and mechanisms are scale invariant, focus should be on how scales relevant to the organisms affect distribution patterns. However, in polar regions where glaciers or a persistence ice foot covers the intertidal no communities are present [35,65], and where the intertidal is characterized by a seasonal ice-foot formation, colonization is only possible during spring and summer [66]. Any future reduction of glaciers, ice shelves and ice foots in these regions will therefore permit organisms to colonize the intertidal zone [67–70]. This will result in increased biomass and α-diversity as is evident from Svalbard in the high Arctic where recent change has resulted in just this outcome [38,71]. Given that most studies fail to consider the aspects discussed above, and that data available from tropical and polar regions are sparse or absent, the relative impacts of multiple stressors ranging from local-to large-scales cannot be explored in depth for intertidal biodiversity. However, our data show that latitudinal patterns in intertidal biodiversity differ significantly from subtidal studies. They also highlight the need for re-evaluating conservation efforts and climate change predictions for this unique global ecosystem.

### Latitudinal gradients in functional groups

In contrast to overall α-diversity, we identified strong latitudinal patterns in assembly composition. The relative dominance of the four functional groups changed with latitudes with predators displaying the strongest gradient. In the northern hemisphere, the number of predators decline at latitudes above 60°N, supporting local-scale studies showing that although intertidal predators, such as crabs and starfish, are found on mid-latitude intertidal shores [72,73], they are missing or rare on exposed Arctic shorelines [36,38,74]. A latitudinal reduction in the number of predators is consistent with the hypothesis of a general reduced level of predation in the polar regions [35,42,75], and similar latitudinal predation clines are found in subtidal benthic habitats [20,21]. Several explanations have been proposed for this latitudinal decline, including the lack of durophagous predators in Antarctica, which has been attributed to the increased solubility of calcium carbonate (CaCO3) with latitude, increasing the cost of shell production [16,76]. The power muscles generate is also strongly affected by temperature and has been proposed as another reason why durophages are absent in high polar regions [35,75], but there is still no full consensus on the processes behind these patterns, and the mechanisms explaining predation gradients remain debated. Predators are important for organizing food webs and community composition. For example, the habitat-forming mussel *(Mytilus californianus)* expanded and excluded more than 25 species after removal of its main predator, the starfish *Pisaster,* in Western North America [73], and in the absence of predators, a tropical seagrass community can support 10 times more species than with predators present [77]. Thus, the low proportion of predators at high latitudes indicates, that the importance of top-down biological interactions in assemblages decreases with latitude in marine ecosystems, and that species composition in polar regions may primarily be controlled by the physical environment [35,36,42,78]. However, it should be noted that we only studied benthic intertidal predators collected during low tides. Therefore, we cannot estimate impacts of mobile predators (e.g. fish and birds) or the predation pressure from epibenthic species during submersion of the shoreline, which can be significant [79,80]. Algae formed the most abundant functional group at high latitudes, consisting of both encrusting species surviving in surface depressions, protected from ice scour and extreme temperatures, and larger macrophytes. Some macrophytes found in polar regions are able to vegetatively regenerate tissue after substantial ice damage to fronds and fastholds, and can survive extreme sub-zero air temperatures during emersion [81]. Indeed air temperature is the prime stressor in polar intertidal systems [82,83], and microhabitats created by macroalgal fronds are important on intertidal shores across latitudes as they provide protection that produces microclimates permitting survival during temperature extremes. The highest latitude shores contain fewer visually obvious species [60], but coralline algae and encrusting species are capable of surviving extreme temperatures in such microhabitats (and in crevices or between boulders), and can survive the winter underneath the intertidal ice foot, likely in protected air pockets or brine pools reached occasionally and/or tidally by water [37,84].

In this study grazers occurred across all latitudes. Grazing can have powerful effects on algal biomass and distribution at low-to mid-latitudes [72,85], and grazing is an important driver of microalgal community structure on King George Island in Antarctica [86]. Contrary to this, no patellid limpets has been reported from south Greenland, suggesting grazing is of little importance in this sub-Arctic region[36]. Quantitative data on grazing impacts in polar intertidal habitats are currently too limited to make conclusions on the ecological implications and effects on latitudinal gradients, yet the inverse latitudinal gradient in algal diversity points to a potential shift in the importance of biological interactions from being severely competitive at lower latitudes to facilitative at high latitudes [42,87,88]. However, to understand ecological implications of latitudinal changes in functional diversity, data are insufficient on the magnitude and influence of biotic interactions across multiple localities from polar to tropical regions. Such research is needed to identify the factors that determine the latitudinal changes in functional diversity identified here, and hence their ecosystem-wide cascading effects.

## Materials and Methods

### Data collection

In 2019 we searched the literature using Web of Science^®^, Google Scholar^®^, and Scopus^®^ for publications devoted to rocky intertidal local species richness, community composition or α-diversity across all latitudes from the Antarctic to Arctic. We only included data from mid-intertidal studies to eliminate confounding effects of intertidal vertical position and tidal amplitude. Reports were considered valid when authors presented assembly-wide data on “α-diversity”, “richness” or “number of species” in terms of species lists, tables or graphs. Specifically, we excluded studies that focused exclusively on algae [26,49], invertebrates [50,51] or individual taxonomic groups [e.g. 24,25,51-53,87]. Full species lists were extracted whenever possible, and WebPlotDigitizer was used to extract graphic data [90]. Additional data were obtained from the Multi-Agency Rocky Intertidal Network (pacificrockyintertidal.org), the Alaska Ocean Observing System (aoos.org), and the South American Research Group on Coastal Ecosystems (SARCE) via the Ocean Biogeographic Information System (obis.org) website. The supplementary Appendix 1 – data sources provides a complete literature list, and is available from the Zenodo repository [91]

In any of the sources used in this study to abstract data, the fraction of species sampled at a given site (inventory completeness) depends on the sampling effort by the collector and local conditions. There are statistical methods to minimize the influence of sampling biases, such as the jackknife estimator of species richness[92], which has been developed to estimate regional γ-diversity from subsamples and abundance [28,93]. However, we focused on α-diversity obtained from published records, tables and figures, and these data were often presented as a single value without supplementary information on subsamples or abundances. Thus, we were unable to verify and compare sampling efforts within sites, but instead we collected data from 433 sites to give very large spatial resolution. To further minimize sampling bias among sites, we only considered studies where all species were collected according to described proscribed set of methodologies.

We validated taxonomic status by checking all the names in the World Register of Marine Species (WoRMS; http://www.marinespecies.org). Whenever species lists were available, we allocated species to functional groups (Fig. 1); Species were allocated to one of four functional groups based on food acquisition; algae (all primary producers), grazers (including scrapers), predators (including scavengers) and suspension-feeders. The final the distribution of algae, grazers, predators and suspension-feeders were analysed and presented as the relative percentage of all species found in that study.

### Statistical analysis

Generalized linear mixed models (GLMMs) with a Poisson distribution were used to model correlations between α-diversity and latitude, and six latitudinal-scale oceanographic drivers (chlorophyll *a*, nitrate, phosphate, salinity, sea surface temperature and silicate) available from NASA Earth Observations (NEO) and USA National Ocean and Atmospheric Administration (NOAA) World Ocean Atlas as means across 5° latitudinal bands. Sites across all longitudes was binned within corresponding 5° latitudinal bands, which was used as a random factor to account for dependency among sites nested within bands.

Latitudinal patterns were analysed separately for the northern and southern hemispheres. Variograms (using the R package gstat [94]) showed spatial autocorrelation among sites from adjacent coastlines, and a variance inflation factor (VIF) analysis showed high levels of collinearity among latitude and temperature (collinearity threshold of 10 [95]). Correlations among the six oceanographic covariates was also assed using VIF, but showed no collinearity (VIF <10). Consequently, correlations between α-diversity and latitude were analysed separately from correlations between α-diversity and oceanographic covariates, to avoid false estimates [96]. To deal with spatial autocorrelation, all GLMMs were fitted with a Matérn spatial covariance structure with longitudinal and latitudinal values as spatial variables [97,98], using the spaMM R package [99]. Model selection showed no significant interactions among oceanographic covariates on α-diversity patterns, and the final full models were built without interactions. The final models were validated by plotting standardized residual patterns plotted against fitted values, and using residual diagnostic tools from the DHARMa R package [100].

To investigate the general influence of local environmental stressors (stressors relevant to species at site-level), data on four local-scale environmental drivers (ice-scour, macroalgal cover, salinity and wave exposure) were extracted from the published literature (see method described above) when available through text, tables or direct author correspondence. This data extraction resulted in four individual datasets, which were analysed separately as most papers only investigated one of the oceanographic drivers; Effects of salinity and wave exposure were separately analysed using fitted standardized input variables (scale from 0 to 1) to estimate effect sizes of data originating from different scales, depending on the literature source [101]. We analysed salinity effects using a GLM with a Poisson distribution, while ice scour, and macroalgal cover effects were analysed using a GLM with a negative binomial distribution because α-diversity was characterised by over-dispersion (determined after visual inspection of residual patterns), but showed no zero-inflation [102]. Wave exposure effects were initially analysed using a GLM, but model residuals showed a non-linear pattern, and the data were re-analysed using a generalized additive model (GAM) with a negative binomial distribution to ensure acceptable residual patterns and normality [103].

All models were finally validated by inspection of standardized residual patterns plotted against fitted values [104] and the proportion of variance explained was calculated as R^2^ =1 – (model deviance/null deviance).

The use of p-values remains debated for GLMMs [105], and we present 95% confidence intervals (95% CIs) for the regression parameters and effect size, estimated using bias-corrected parametric bootstrap methods (10,000 iterations) [104]. The effect of the variable was considered significant when 95% CIs did not overlap zero.

All statistical analysis and data exploration were conducted using R [106]. Initial data exploration, data analysis and presentation followed [96,104].

## Acknowledgements

The authors thanks Brenda Konar, Takashi Noda, Tan Koh Siang, Robin Elahi, Valdivia Nelson and Megan Dethier for providing intertidal data. We gratefully acknowledge the Valdivia Nelson’s projects FONDECYT (grant #1141037) and CONICYT-PIA (ANILLO; grant #ART1101) for providing data from Chile and Antarctic, respectively. This study further utilized data collected by the South American Research Group on Coastal Ecosystems (SARCE) sponsored by TOTAL foundation, and data collected by the Multi-Agency Rocky Intertidal Network (MARINe): a longterm ecological consortium funded and supported by many groups. Please visit pacificrockyintertidal.org for a complete list of the MARINe partners responsible for monitoring and funding these data. Data management has been primarily supported by BOEM (Bureau of Ocean Energy Management), NPS (National Parks Service), The David & Lucile Packard Foundation, and United States Navy.’ This work is a contribution to ‘The future of Arctic biodiversity in a climate change era’ project.

## Data availability statement

Data used in this study is freely available from the Multi-Agency Rocky Intertidal Network (MARINe; pacificrockyintertidal.org), the Ocean Biogeographic Information System (OBIS; obis.org) and the Alaska Ocean Observing System (AOOS; aoos.org). Publications obtained from the literature search, and previously unpublished species lists from Arctic locations can be found on the Zenodo data repository [91].

